# Millisecond Mix-and-Quench Crystallography (MMQX) Enables Time-Resolved Studies of PEPCK With Remote Data Collection

**DOI:** 10.1101/2021.04.06.438620

**Authors:** Jonathan A. Clinger, David W. Moreau, Matthew J. McLeod, Todd Holyoak, Robert E. Thorne

## Abstract

Time-resolved crystallography of biomolecules in action has advanced rapidly as methods for serial crystallography have improved, but the large number of crystals and complex experimental infrastructure required remain serious obstacles to widespread application. We have developed Millisecond Mix-and-Quench crystallography (MMQX), which yields millisecond time-resolved data using far fewer crystals and routine remote synchrotron data collection. To demonstrate the capabilities of MMQX, the conversion of oxaloacetic acid to phosphoenolpyruvate by phosphoenolpyruvate carboxykinase is observed with a time resolution of 40 ms. MMQX, by lowering the entry barrier to time-resolved crystallography, should enable broad expansion in structural studies of protein dynamics.

## Introduction

Extending X-ray crystallography to yield ensemble-averaged all-atom, atomic-resolution molecular movies of enzymes in action has long been a major goal in structural biology.^1–6^ Early time-resolved crystallography (TRX) experiments examined reactions that reset within the crystal and were triggered optically, such as CO release by myoglobin and the photocycle of photoactive yellow protein (PYP).^2,3^ These features allowed complete data sets at multiple time points to be obtained using a small number of crystals, with radiation damage limiting the amount of data collected from each crystal. The development of X-ray free electron lasers (XFELs) - with ultrabright femtosecond X-ray pulses that enable capture of single-frame, fixed-crystal orientation snapshots before most damage occurs - and of serial microcrystallography - enabling delivery of tens of thousands of randomly oriented microcrystals into the XFEL beam and analysis of the resulting diffraction data sets - have driven major advances in TRX methods.^7–11^ These have been adapted for microbeam synchrotron sources.^12–15^ Microcrystals enable chemical triggering via diffusion on millisecond timescales, short enough to reveal biologically important intermediate states.

Optical excitation and/or crystal mixing and delivery systems are complex and must be integrated into the X-ray beamline for TRX experiments. Optimization for a given protein crystal system and efficient serial data collection requires multiple collaborators, knowledgeable beamline staff, and often extended beamline access. Complex sample delivery systems, large sample consumption, and limited access to suitably-configured beamlines are formidable obstacles to broad application of TRX techniques.

Freeze-quenching techniques provide an alternative approach to obtaining time resolved data that have been productively used in spectroscopic studies (e.g., EPR) of liquid and powder samples^16,17^ and, more recently, in single-particle cryo-electron microscopy analysis of ribosomes to observe conformational changes during elongation factor binding.^18,19^ Unlike in time-resolved studies conducted at room/biological temperature, freeze-quenching allows the reaction and measurement times to be separated, and long measurement times allow more data to be collected per sample per time point. Furthermore, in crystallography, radiation damage per unit dose is dramatically reduced at cryogenic temperatures, allowing more data to be collected per crystal.

Freeze-quenching was not considered viable for TRX studies, largely because reactant diffusion and crystal cooling times for crystals large enough to yield adequate diffraction were longer than most timescales of biological interest.^20–23^ However, the high brilliance of both XFELs and current generation storage ring light sources now enable data collection from micrometer size crystals, for which diffusion times can be below 5 ms (Supplemental Table S2). Advances in cryocooling technology based on high speed plunging and removal of cold gas layers allow these crystals to be cooled to below 150 K in <3 ms.^22^ Diffusion and cooling times on this scale should allow observation of slow dynamic processes and long-lived chemical intermediates.^23^

We have developed a technique for mix-and-quench time-resolved crystallography by introducing substrate into microcrystals during a robotic plunge into liquid nitrogen (LN_2_). This technique generates millisecond time-resolved structures using one or a few crystals per time point in a single shift of standard remote synchrotron data collection. MMQX thus promises to make TRX data collection much more broadly accessible.

## Results

In order to initiate biochemical reactions in crystals, we developed a plunge-through-film approach (Figure 1). A film containing substrate that spans a wire loop is placed in the plunge path between the crystal’s initial position and the LN_2_ reservoir. To create a suitably large and stable film in the loop and to cryoprotect the crystal, the film contains 3% w/v SDS and 20% v/v PEG400, respectively. The crystal on its mount is robotically plunged through the loop at 2 m/s, piercing the film which then collapses around all sides of the crystal and the polyimide MicroGripper sample holder. To ensure that liquid from the film immediately contacts all sides of the crystal’s surface and begins diffusing to its interior, excess liquid surrounding the crystal is blotted away in an 85% r.h. environment right before the plunge. To ensure that the reaction proceeds at room temperature until quenching and to maximize crystal cooling rates, cold gas that would normally be present to a height of 2-6 cm above the LN_2_ surface^22^ is carefully removed using a flow of dry room temperature N_2_ gas and matched suction (Figure 1), giving a transition from T>273 K N_2_ gas to 77 K LN_2_ over a distance of ∼0.1 mm (Supplemental Figure 1A). Cooling times from 273 K to 150 K of ∼6 ms were measured using 50 μm thick thermocouples (Supplemental Figure 1B).

**Fig. 1.**
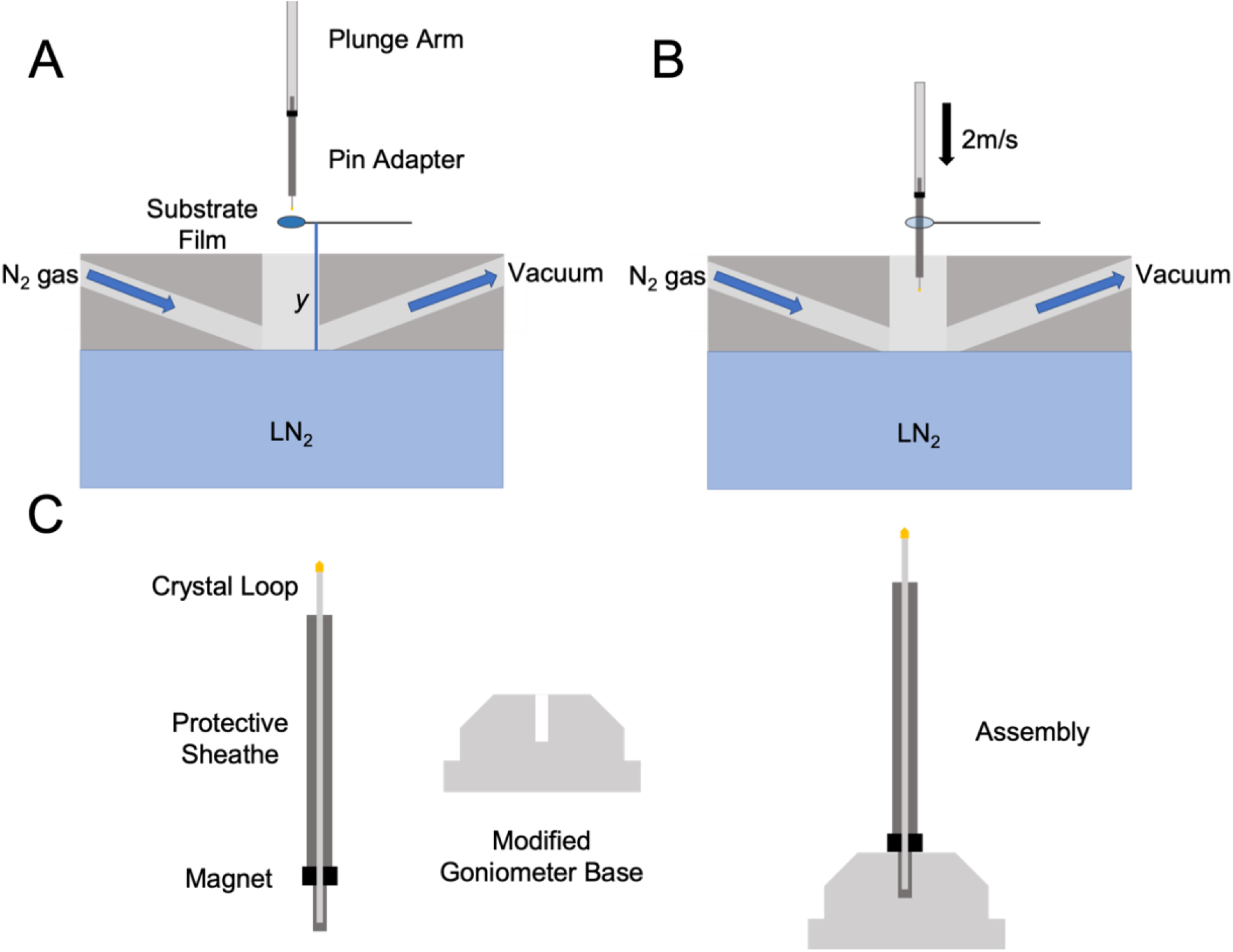
**A**, Plunge cooler design for MMQX, including a substrate-containing film. A polymer crystal mount on a stainless steel pin in a magnetic mount holder is placed on a plunge arm attached to a high-speed linear translation stage. To remove cold N_2_ gas that forms above the LN_2_, warm N_2_ gas flows in through a manifold above the LN_2_ surface and then is removed via suction. Time point is determined by height of film above LN_2_ (*y*) and plunge rate. **B**, A crystal is plunged through the substrate film at a speed of 2 m/s and then through room temperature N_2_ gas and through the LN_2_ surface. **C**, Crystal pin holder for rapid plunge cooling and MMQX experiments. To reduce the aperture size of the substrate-containing loop, the crystal is plunged in the pin holder. Once cooled, the pins are inserted into modified ALS-format goniometer bases.

To test this approach, we used crystals of phosphoenolpyruvate carboxykinase (PEPCK), an essential enzyme in glucose metabolism that catalyzes the reversible, nucleotide-dependent interconversion of oxaloacetic acid (OAA) and phosphoenolpyruvate (PEP) using Mn^2+^ as a metal cofactor.^24^ The kinetics and mechanism of this reaction are well understood, starting with OAA decarboxylation to form an enol-pyruvate intermediate, which then attacks the γ-phosphate of GTP.^25^ Structures obtained by co-crystallization and soaking have examined binding of ligands including GTP, GDP, Mn^2+^, OAA, and PEP. In PEP-containing structures deposited in the PDB, PEP is not found in the catalytically competent phosphate transfer site, but at a secondary site referred to as the outer-shell product binding pocket, thought to be PEP’s locale prior to release. We used rod-like Mn^2+^ GTP-PEPCK co-crystals with dimensions of 100 μm × 20 μm × 8 μm, and a film containing OAA. With these crystals and our apparatus, Δ*t*_*plunge*_= 0.5 ms per mm of height Δ*y*; Δ*t*_*cool*_ was conservatively estimated to be <10 ms; and Δ*t*_*diffuse*_ is ≈12 ms (Supplemental Table 2).

In electron density maps derived from diffraction data sets corresponding to Δ*t* ≈ 40 ms, we observe the PEP product in the phosphate transfer site, complexed by the Mn^2+^ active site ion, not in the product pocket as in previous PEP structures (Figure 2B and Figure 3). This previously unobserved PEP position is the expected position immediately following phosphate transfer from GTP. The phosphate-to-phosphate distance of 3.4 Å compares with 7.5 Å in previous PEP+GDP structures. Our 120 ms time point also shows PEP in the phosphate transfer site, not the product binding pocket, suggesting that in the presence of nucleotide, movement to the release site position is slow relative to PEP release (Figure 2C). Our 40ms and 120ms post-mixing structures also contain CO_2_, which was formed from OAA during the generation of the enol-pyruvate intermediate (Figure 3). The chemical identity of the carboxylating agent as well as its specific binding site have been contentious^26,27^ and this TRX data agrees with prior kinetic data indicating that CO_2_ (rather than bicarbonate) is the form of the substrate for carboxylation of PEP.^28^ Our 120 ms structure shows relaxation relative to the 40 ms structure of CO_2_ to a different orientation in the binding pocket, where it has favorable interactions with the main chain nitrogens of Arg87 and Gly237, as well as the Asn403 sidechain.

**Fig. 2.**
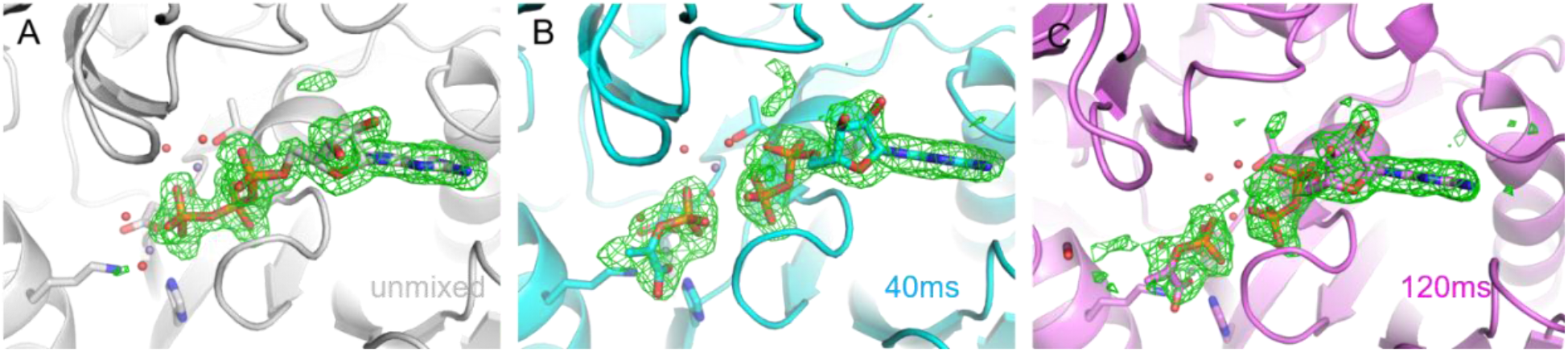
MMQX experiments show diffusion of OAA into the PEPCK active site and formation of PEP product. **A**, Polder map of unmixed PEPCK (gray) data omitting GTP. **B**, 40 ms time point (cyan) Polder map omitting PEP and GDP. **C**, 120 ms time point (magenta) Polder map omitting PEP and GDP. All Polder maps are shown at 4 RMSD.

**Fig. 3.**
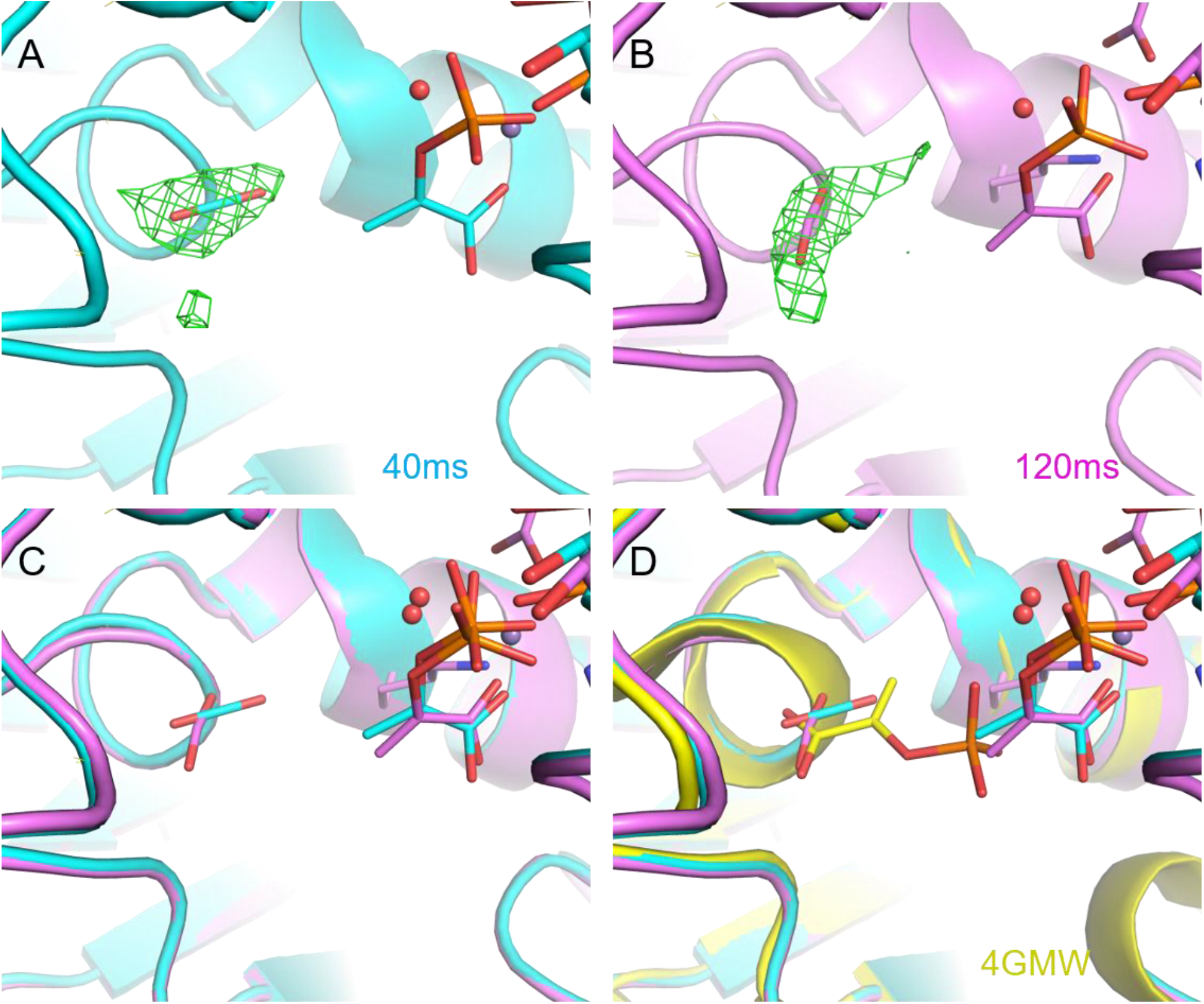
Comparison of CO_2_ binding sites at 40 ms and 120 ms, and with a previous PEP-containing PEPCK structure. **A**, Polder map at 4.5 RMSD of CO_2_ from the 40 ms MMQX data. **B**, Polder map at 4.5 RMSD of CO_2_ from the 120 ms MMQX data. **C**, Comparison of the CO_2_ binding site at these two time points. D, Representative location of PEP from PDB ID 4gmw compared with the 40 ms and 120 ms time points.

As the unmixed and Δ*t* ≈ 40 ms data are isomorphous, maps corresponding to F_O mixed_-F_O unmixed_ can be calculated. These show clear hallmarks of GTP to GDP conversion, with relaxation of the α− and β−phosphates away from the Mn^2+^ catalytic ions (Figure 4A and 4B). The surrounding residues follow the GDP relaxation and breathe into the GDP binding pocket (Supplemental Figure 2. As the Δ*t* ≈120 ms data is no longer isomorphous with the unmixed data, we cannot use as sensitive a method for analysis, but general inspection of the data suggests that little has changed in the active site between the 40 and 120 ms time points. However, density further away from the active site is noticeably less well defined at 120 ms than in either the 40 ms or unmixed data, contributing to worse refinement statistics. This may be due to crystal-to-crystal variation or to partially constrained motions involved in product release. The Ω gating loop in all our structures is open and disordered, consistent with the hypothesis that the loop is open in unoccupied active sites as well as during the product release steps after the reaction is catalyzed.^25^

**Fig. 4.**
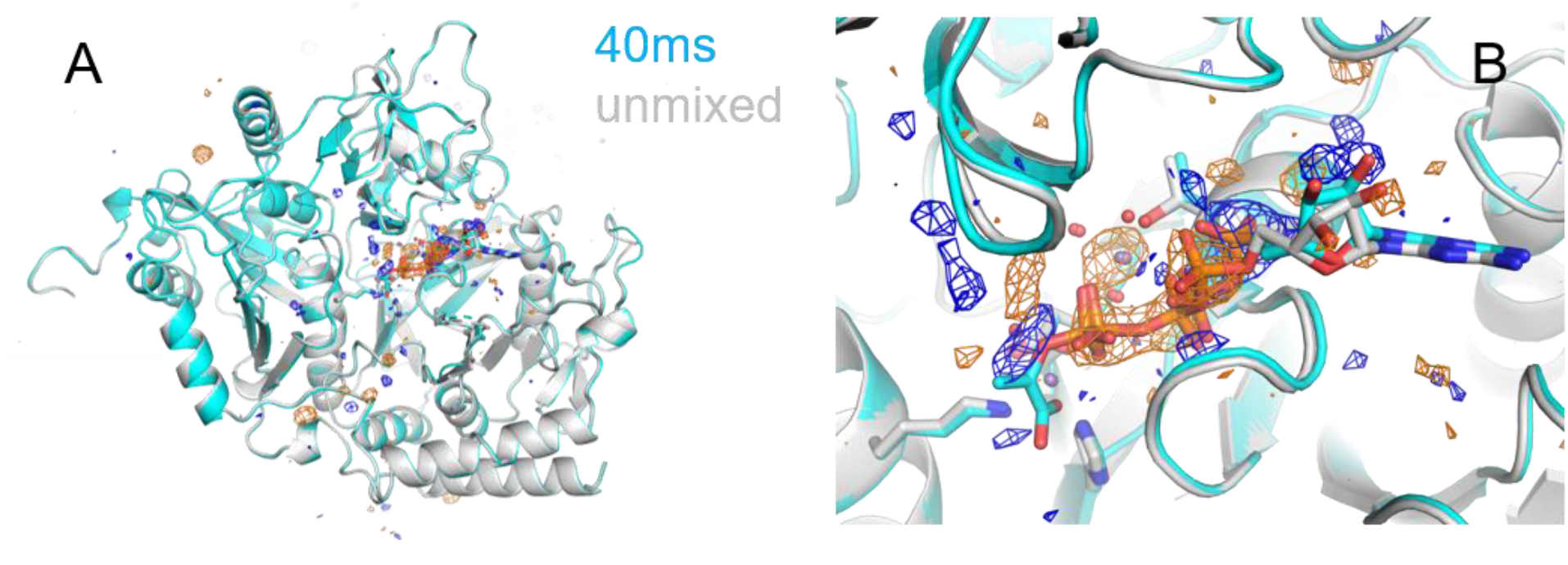
FO-FO analysis of 40 ms (cyan)-unmixed (gray) PEPCK data. FO-FO maps shown at ±4 RMSD (blue, orange). **A**, Overview of the asymmetric unit demonstrating that large difference map peaks are concentrated in the active site. **B**, View of active site showing difference map peaks for PEP and relaxation of GDP.

## Discussion

We have demonstrated the feasibility of millisecond mix-and-quench crystallography, MMQX, using it to trap a previously unobserved structural state of an enzyme *in crystallo*. This method can yield time resolutions of a few ms (Supplemental Section 2) and is generalizable to many enzymatic systems. Its sample usage is comparable to that in standard cryo-crystallography and at least two orders of magnitude less than in the most sample-efficient room-temperature synchrotron-based time-resolved methods, and achieves time resolution comparable to the best achieved using serial mixing crystallography.^9,29^ Millisecond time-resolved information can be obtained using mail-in remote data collection on standard cryo-crystallography synchrotron beamlines without modification, and using crystals left over from initial structure determinations. Consequently, this method should expand access to the high-impact field of time-resolved crystallography and enable structural enzymology to become a powerful tool for many scientists. MMQX is complementary to time-resolved serial femtosecond crystallographic methods. Data collection under physiological conditions with negligible radiation damage is exchanged for orders of magnitude less sample consumption, much simpler sample preparation, and much greater beamtime access for iteration and optimization.

## Methods

### Plunge Cooler Design

The plunge cooler was based on a prototype design provided by MiTeGen, LLC, that we heavily modified and adapted for our purposes. As shown in Supplemental Fig. 1, the cooler consists of a vertical translation stage for the sample, a gas management manifold immediately above the liquid nitrogen (LN_2_) having a central bore coaxial with the plunge path, and an insulated chamber containing the LN_2_. The vertical translation stage has a carriage to which the sample arm is attached, and this carriage moves up and down on a lead screw driven by a DC motor. The carriage accelerates from rest to a maximum speed of 2 m/s, which it then maintains until it reaches a soft stop at a position where the sample is several centimeters below the LN_2_ surface.

The gas management manifold has ports for dry room temperature N_2_ gas and for suction provided by a Venturi vacuum generator. Heaters and insulation keep manifold surfaces above the LN_2_ near room temperature. Prior to plunging, cold gas that forms above the LN_2_ is removed using warm dry N_2_ gas that is projected in a laminar flow across the LN_2_ surface and is removed via the suction port, with both gas and suction controlled via computer using solenoid valves. As shown in Supplemental Fig. 2A, with this cold gas removal system and other design features of the manifold, the temperature remains above 273 K to within 100 μm of the LN_2_ surface.^22^

Supplemental Figure 2B shows temperature versus time during a plunge into the LN_2_ recorded using a thermocouple with a bead roughly 100 μm in diameter and 50 μm thick. The time for the thermocouple to cool from the freezing point of pure water, 273 K, to the protein-solvent glass transition near 200 K is ∼3.5 ms, and to the glass transition temperature of pure water T_g_∼136 K is ∼6 ms. These time intervals correspond to average cooling rates of ∼20,000 K/s. Thermocouple metals have nearly the same heat capacity per unit volume between 273 K and 200 K or 136 K as protein crystals (including the latent heat of fusion of internal water). In this size range cooling rates are determined by the thermal boundary layer in the surrounding N_2_, not the thermal conductivity of the sample. Consequently, thermocouples and protein crystals of similar size should cool at similar rates. For rod-shaped PEPCK crystals with transverse dimensions of 8 μm × 20 μm, visual observation of frozen samples suggested a thickness of crystal + (substrate and PEG-containing) solution of roughly 20 μm, so cooling rates were likely >60,000 K/s^21^ and cooling times from 273 K to 200 K <1.2 ms. With crystals a few micrometers in size and total crystal + liquid thicknesses of <10 μm, cooling rates in excess of 100,000 K/s should be achievable, corresponding to cooling times to 200 K of ∼0.7 ms.

### Sample Preparation

PEPCK crystals were prepared as previously described.^30^ Briefly, rat cytosolic PEPCK protein was expressed, purified, and concentrated to 10 mg/mL for crystallization experiments. PEPCK solution containing 20 mM MnCl_2_ and 5 mM GTP (both added immediately prior to crystallization) was mixed 1μL:1μL in sitting drop crystallization plates with reservoir solution containing 100mM HEPES pH 7.5 and 15-20% w/v PEG3350. Suitable crystals were recovered from wells containing 19% w/v PEG3350. The crystals used were rod-like with dimensions of 100 μm × 20 μm × 8 μm.

### Mixing Experiment

PEPCK crystals were supported during plunging and data collection on MiTeGen MicroGrippers™, which have narrow fingers projecting radially inward from an outer loop that support the crystal while maximizing unobstructed access to the crystal surface. Crystal harvesting and placement on the plunge cooler’s sample carriage was performed within an ∼85% r.h. environment to minimize crystal dehydration between harvest and plunge. As shown in Figure 1A, a thin substrate-containing film spanning a circular wire loop was placed along the crystal’s plunge path at a measured height above the LN2 surface. For PEPCK, these films contained 20 mM OAA, 20% v/v PEG 400, and 3% w/v SDS. OAA was the substrate in the reaction, PEG 400 provided cryoprotection of solution transferred from the film to the crystal surface, and SDS increased the film’s stability. The stainless steel pin of the MicroGripper was inserted into a cylindrical pin adapter of a custom two-part goniometer base (Figure 1C); this part had a diameter sized to pass through the film-containing loop, and was attached to the sample arm on the sample carriage with a similar diameter stainless steel rod. Crystals were then robotically plunged at 2 m/s through the film, which was positioned at varying heights above the LN_2_ surface, creating different delays between the start of substrate diffusion into the crystal and the start of cryocooling. The cold gas layer that would normally be present above the LN_2_ was efficiently removed so that the gas temperature remained above 273 K to within 100 μm of the LN_2_ surface (Supplemental Fig. 1A), ensuring that substrate diffusion and reaction was not slowed by cooling until the crystal entered the LN_2_. After cooling/quenching, the sample + the pin adapter was removed from the plunge cooler’s sample arm and inserted into part 2 of the goniometer base. The assembly was then loaded into UniPucks for later data collection.

The time point Δ*t* of the resulting mix-and-quench dataset is estimated as Δ*t* = Δ*t*_*plunge*_ + Δ*t*_*cool*_, where Δ*t*_*plunge*_ = *v*_*plunge*_Δ*y* is the time for the crystal to plunge the distance Δ*y* between the substrate-containing film and the LN_2_ surface, Δ*t*_*cool*_ is the time for the sample to cool between room temperature and the protein-solvent glass transition near 200 K; and Δ*t*_*diffuse*_ is the time for substrate to diffuse into the crystal, averaged over the X-ray illuminated crystal volume. With Δ*t*_*cool*_ < 10 ms, Δ*t*_*diffuse*_ = 12 ms, and *v*_*plunge*_ = 2 m/s, Δ*y* values of 60 mm, and 220 mm correspond to Δ*t*_*plunge*_values of 30 and 110 ms and Δ*t* values of 40 and 120 ms. The smallest feasible distance Δ*y* (limited by radiative cooling of the substrate-containing film, which can be minimized by translating the film into place immediately before the crystal plunge)– is likely of order 4 mm, corresponding to Δ*t*_*plunge*_ ∼2 ms.

### Data Collection and Processing

PEPCK diffraction data was collected at the CHESS FLEXX beamline (Beamline 7B2) using a CRL lens-focused 9 μm × 12 μm x-ray beam containing 2 × 10^11^ ph/s at an energy of 11 keV. The sample temperature was maintained at 100 K using a N_2_ gas cryostream, and diffraction data recorded using fine slicing on a Pilatus 6M detector. Raw diffraction frames were processed with DIALS.^31^ Data scaling and merging was performed with aimless.^32^ Phenix.refine and Coot were used for model building and refinement.^33,34^ FO-FO maps using substrate-mixed data at a nominal Δ*t* ≈ 40 ms and unmixed data were carried out to the maximum resolution of the 40 ms data set of 2.17Å, using the well-refined unmixed structure at 1.84Å for phase information. POLDER-OMIT maps were calculated from each data set while omitting GTP in the unmixed data and either PEP, carbon dioxide, or GDP.^35^ See Supplemental Table 1 for diffraction and refinement statistics.

## Acknowledgements

The authors would like to thank George N. Phillips Jr. and Mitch Miller for productive conversations and advice in adapting time-resolved crystallography to standard synchrotron sources. This work is based upon research conducted at the Cornell High Energy Synchrotron Source (CHESS), which is supported by the National Science Foundation and the National Institutes of Health/National Institute of General Medical Sciences under NSF award DMR-1829070, using the Macromolecular Diffraction at CHESS (MacCHESS) facility, which is supported by award GM-124166 from the National Institutes of Health, through its National Institute of General Medical Sciences. JAC, DWM, MJM, and RET were supported by award GM127528-02 from the National Institutes of Health, through its National Institute of General Medical Sciences. TH and MJM acknowledge support from the Natural Sciences and Engineering Research Council of Canada.

## Author Contributions

JAC conducted the experiments with support from DWM. MJM expressed and purified the PEPCK samples. JAC prepared the PEPCK crystals. JAC processed and analyzed the diffraction data with assistance from MJM and TH. DWM and RET designed the plunge cooler system. JAC and DWM adapted the plunge cooler for time-resolved studies. The manuscript was written by JAC, DWM, and RET with contributions from all authors.

## Online Content

Further information pertaining to the method, additional references, supplemental information, and diffraction statistics are available at DOI:

