## Supplemental Material for "Millisecond Mix-and-Quench Crystallography (MMQX) Enables Time-Resolved Studies of PEPCK With Remote Data Collection"

### 1. Thermocouple measurements characterizing the performance of the plunge cooler

Supplemental Figure 2A shows the temperature, measured using a type-K thermocouple having a bead roughly 100 μm in diameter and 50 μm thick, as a function of height *y* above the surface of the liquid $N_2$ in the plunge cooler's insulating container. The thermocouple was slowly stepped toward the surface, and warm dry $N_2$ gas and suction applied using the plunge cooler's gas management manifold to remove cold gas that formed above the liquid nitrogen. Supplemental Figure 2B shows the temperature versus time measured for different plunge speeds when using the same thermocouple as in 2A and the gas management manifold to remove cold gas that formed above the $LN_2$. The thermal mass and cooling time of the thermocouple are expected to be larger than those of the 100 μm × 20 μm × 8 μm crystals used here.

### 2. Estimates of diffusion times for OAA and glucose

Our estimates of diffusion times for OAA into PEPCK crystals and glucose into GI crystals – are based on a solution to Fick's second law for a cuboid crystal described by Schmidt.[1,2] This analysis ignores the fact that the solvent is nanoconfined within channels of a protein crystal, that viscosity may be affected by this confinement, and that substrate may have weak binding interactions with the protein surface, all of which are likely to slow diffusion.[3,4] This analysis does not account for any solvent present on the crystal surface prior to mixing, which could slow diffusion by providing additional liquid through which the substrate must diffuse and by reducing the substrate concentration at the crystal surface.[2] Here, considerable care was taken to remove that solvent. No diffusion constant measurements for OAA are available. The diffusion constant of erythritol, which has a similar molecular weight and hydrodynamic radius to OAA, of $7.6 \times 10^{-6}$ $cm^2/s$ in dilute aqueous solution at 295 K[5] is used. Glucose has a diffusion coefficient of $6.3 \times 10^{-6}$ $cm^2/s$ in dilute aqueous solution at 295 K.[6]

Supplemental Table 2 gives the resulting diffusion time estimates for OAA into PEPCK crystals and glucose into glucose isomerase crystals for crystals of different sizes. These estimates involve many assumptions and should be tested in future experiments. Note that any precooling of the sample in cold gas as it descends from the substrate-containing loop to the $LN_2$ surface – which is avoided via our plunge cooler design but that would be substantial in ordinary hand-plunging into $LN_2$ contained within foam Dewars – could dramatically increase diffusion times. For example, the diffusion coefficient of glycerol at 273 K is ~40% of its value at 295 K.[7]

**3. Estimate of minimum number of crystals of relevant size for mix-and-quench experiments required to obtain complete data sets at synchrotron sources.**

To estimate the crystal efficiency of MMQX relative to room temperature time-resolved SSX methods, we used the required crystal number / size calculator developed by Holton and Frankel at https://bl831.als.lbl.gov/xtalsize.html (Supplemental Table 2).[8,9] Our calculations make the following assumptions: the crystal is sized to achieve desired mixing time regimes, the X-ray beam size is matched to crystal size, the screening image diffracts to 1.5 Å, and the required merged resolution is 2.0 Å with an $I/\sigma$ at 2.0 Å of 1.4. These parameters are reasonable for our PEPCK MMQX data, as many of our crystals show diffraction peaks to 1.6 Å or higher, and a 2.0 Å cutoff is sufficient to obtain required map features. $I/\sigma$ of 1.4 is between the traditional 2.0 cut-off and the more aggressive merging utilized with $CC_{1/2}$ resolution cutoffs. Our calculations indicate that for PEPCK and glucose isomerase, one crystal should allow collection of a complete dataset to time points of 30 ms or slower, assuming the crystal size matches the time point desired. This increases with macromolecule/asymmetric unit size; for ribosomes at least 13 crystals would be required to collect data at a 30 ms time point for a diffusing ligand comparable in size to glucose. The small numbers of crystals required for MMQX compared with current TR serial crystallography methods allows collection of many more time points with a given sample amount, or experiments to be performed using multiple substrates/ligands. Ongoing advances in data collection procedures, e.g., utilizing X-ray beam offsets[10] or utilizing very hard X-rays[11], at microbeam synchrotron sources will make MMQX more sample efficient.

7. Akinkunmi, F. O., Jahn, D. A. & Giovambattista, N. Effects of temperature on the thermodynamic and dynamical properties of glycerol-water mixtures: A computer

simulation study of three different force fields. *J. Phys. Chem. B* **119**, 6250–6261 (2015).

8. Holton, J. M. A beginner's guide to radiation damage. *J. Synchrotron Radiat.* **16**, 133–142 (2009).

9. Holton, J. M. & Frankel, K. A. The minimum crystal size needed for a complete diffraction data set. *Acta Cryst D.* **66**, 393–408 (2010).

10. Yamamoto, M. *et al.* Protein microcrystallography using synchrotron radiation. *IUCrJ* **4**, 529–539 (2017).

11. Sanishvili, R. *et al.* Radiation damage in protein crystals is reduced with a micron-sized X-ray beam. *Proc. Natl. Acad. Sci. U. S. A.* **108**, 6127–6132 (2011).

**Supplemental Table 1. Data collection and refinement statistics.**

| | Unmixed PEPCK + Mn & GDP | 40ms post OAA mixing PEPCK | 120ms post OAA mixing PEPCK |
|---|---|---|---|
| **PDB ID** | 7L36 | 7L3M | 7L3V |
| **Wavelength** | 1.127 | 1.127 | 1.127 |
| **Resolution range** | 41 - 1.84 (1.906 - 1.84) | 56.88 - 2.17 (2.248 - 2.17) | 41.66 - 1.92 (1.989 - 1.92) |
| **Space group** | P 1 21 1 | P 1 21 1 | P 1 21 1 |
| **Unit cell** | 44.38 118.55 60.23 90 109.53 90 | 44.43 119.01 60.39 90 109.64 90 | 46.12 118.84 61.19 90 107.27 90 |
| **Total reflections** | 470638 (16681) | 348523 (14663) | 432426 (16723) |
| **Unique reflections** | 50261 (4755) | 31241 (3101) | 47723 (4742) |
| **Multiplicity** | 9.4 (3.5) | 11.2 (4.7) | 9.1 (3.5) |
| **Completeness (%)** | 98.84 (93.59) | 99.97 (99.87) | 99.54 (99.35) |
| **Mean I/sigma(I)** | 8.74 (1.17) | 7.33 (2.76) | 7.31 (1.24) |
| **Wilson B-factor** | 15.30 | 6.19 | 5.28 |
| **R-merge** | 0.2284 (1.296) | 0.3262 (0.627) | 0.4032 (1.48) |
| **R-meas** | 0.2421 (1.525) | 0.3415 (0.7035) | 0.4321 (1.733) |
| **R-pim** | 0.07822 (0.7931) | 0.09894 (0.3104) | 0.1484 (0.8891) |
| **CC1/2** | 0.987 (0.381) | 0.975 (0.8) | 0.802 (0.48) |
| **CC*** | 0.997 (0.743) | 0.994 (0.943) | 0.943 (0.805) |
| **Reflections used in refinement** | 50263 (4727) | 31245 (3102) | 47864 (4728) |
| **Reflections used for R-free** | 2610 (245) | 1543 (146) | 2416 (245) |
| **R-work** | 0.1524 (0.2663) | 0.1641 (0.2111) | 0.2023 (0.2918) |
| **R-free** | 0.1988 (0.2992) | 0.2179 (0.2596) | 0.2519 (0.3398) |
| **Number of non-hydrogen atoms** | 5304 | 5234 | 5230 |

| | | | |
|---|---|---|---|
| macromolecules | 4840 | 4865 | 4752 |
| ligands | 36 | 45 | 45 |
| solvent | 428 | 324 | 433 |
| **Protein residues** | 613 | 612 | 601 |
| **RMS(bonds)** | 0.004 | 0.007 | 0.010 |
| **RMS(angles)** | 0.75 | 0.95 | 1.18 |
| **Ramachandran favored (%)** | 97.04 | 97.04 | 96.64 |
| **Ramachandran allowed (%)** | 2.96 | 2.80 | 3.36 |
| **Ramachandran outliers (%)** | 0.00 | 0.16 | 0.00 |
| **Rotamer outliers (%)** | 0.00 | 0.97 | 0.60 |
| **Clashscore** | 3.41 | 5.03 | 5.26 |
| **Average B-factor** | 22.49 | 14.21 | 30.09 |
| macromolecules | 21.94 | 13.75 | 29.74 |
| ligands | 18.77 | 22.20 | 40.45 |
| solvent | 28.99 | 20.12 | 32.89 |
| **Ligand real-space correlation coefficient** | | | |
| GTP | 0.98 | | |
| GDP | | 0.94 | 0.88 |
| PEP | | 0.90 | 0.91 |
| $CO_2$ | | 0.84 | 0.81 |

Statistics for the highest-resolution shell are shown in parentheses.

**Supplemental Table 2:** Estimation of diffusion times and number of crystals required in order to collect a full dataset from crystals that can achieve these mixing times.[8,9]

| Crystal Size ($\mu m^3$) | PEPCK crystals | | Glucose Isomerase Crystals | | Eukaryotic Ribosome crystals |
|---|---|---|---|---|---|
| | OAA diffusion time (ms) | Crystals per dataset | Glucose diffusion time (ms) | Crystals per dataset | Crystals per dataset |
| 2×2×2 | 0.2 | 100 | 0.3 | 240 | 4300 |
| 5×5×5 | 1.1 | 12 | 1.7 | 28 | 490 |
| 10×10×10 | 4.5 | 2 | 6.8 | 4.9 | 85 |
| 10×20×30 | 9.8 | 0.45 | 14.9 | 1.1 | 19 |
| 20×20×20 | 17.8 | 0.31 | 27.1 | 0.75 | 13 |
| 30×30×30 | 40.0 | 0.1 | 60.9 | 0.24 | 4.2 |
| 40×40×40 | 71.2 | 0.04 | 108.2 | 0.11 | 1.8 |

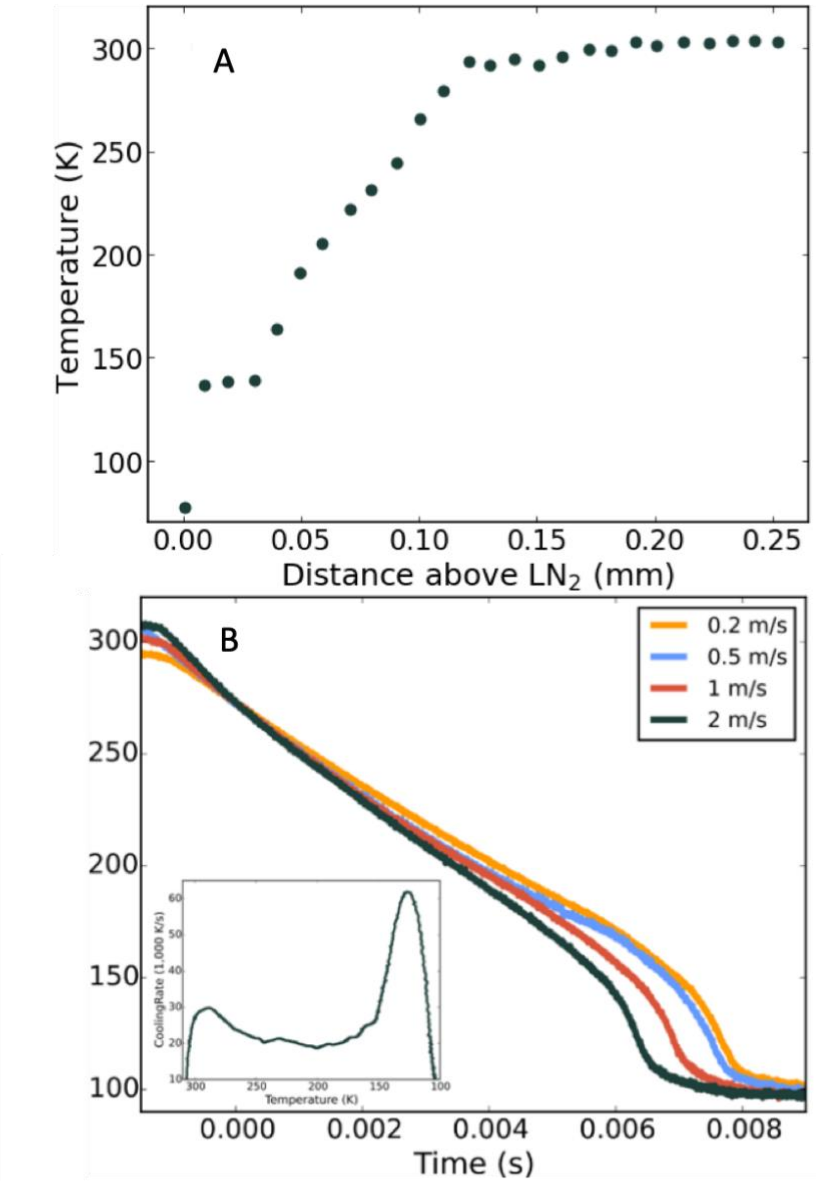


**Supplemental Fig. 1.** Measurements, using a thermocouple with a junction bead roughly 100 μm in diameter and 50 μm thick, characterizing the performance of the plunge cooling system. **A**, Temperature vs height *y* above the $LN_2$ surface along the sample plunge path, measured using a thermocouple held on the sample arm of the plunge cooler. With cold gas removal and replacement by the manifold, the temperature remains above 273 K to within 0.1 mm of the $LN_2$ surface. **B,** Temperature vs time recorded during plunges at speeds between 0.2 and 2 m/s. The time t=0 is set by the time when the thermocouple temperature reaches 273 K, which occurs very close to (just above or below) the $LN_2$ surface. For a plunge speed of 2 m/s, the cooling time from 273 K to 150 K is 5.8 ms. For samples the size of the PEPCK crystals used in our MMQX experiments (rods with a cross section of 20 μm × 8 μm), cooling times from room temperature to 150K should be less than 10 ms and likely on the order of 2 ms.

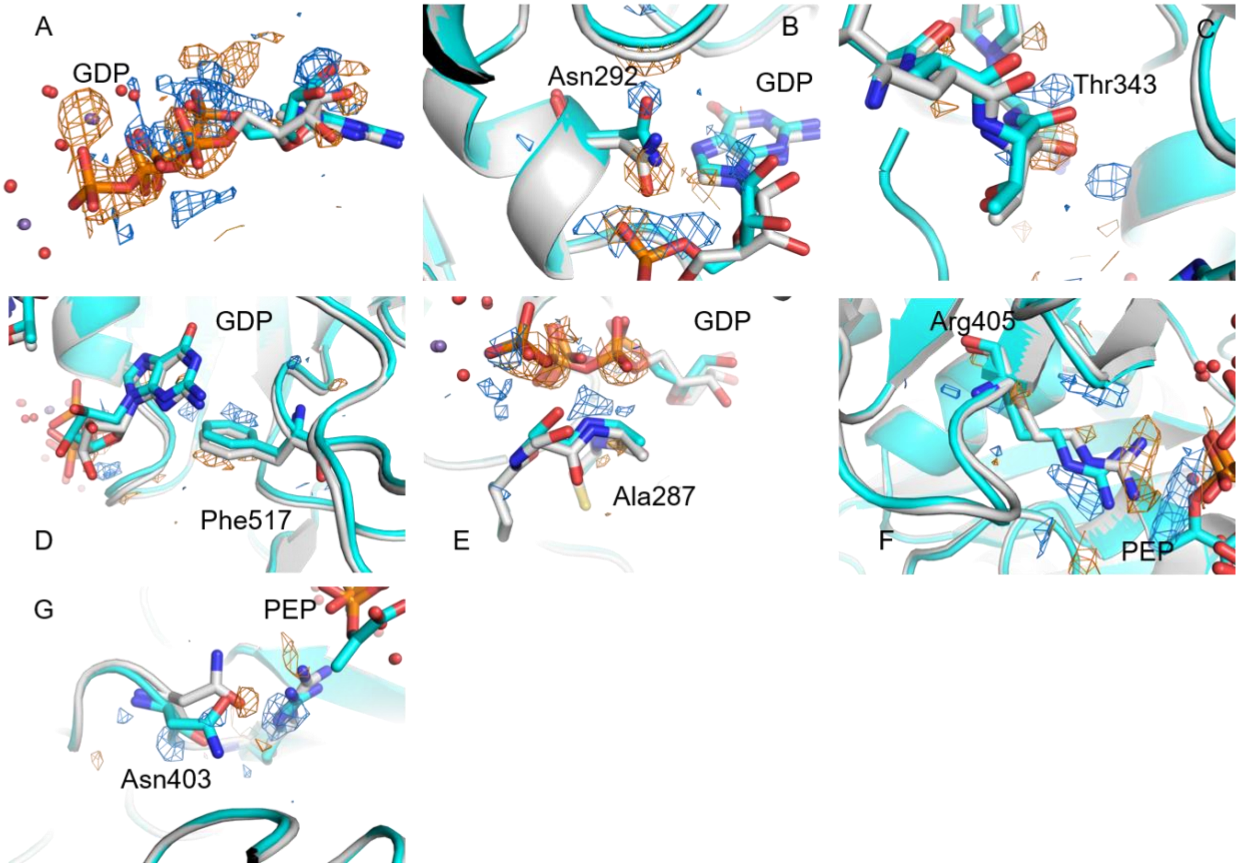


**Supplemental Fig. 2.** Sidechain and GTP binding pocket motions associated with GDP relaxation 40 ms post-reaction. Fo-Fo maps are contoured at ±3.5 RMSD (+blue, -orange). **A**, GTP (gray, umixed)-GDP (cyan, 40ms) relaxation. **B**, Asn292 motion. **C**, Thr343 loop motion. **D**, Phe517 sidechain breathing. **E**, Ala287 motion into phosphate binding cleft. **F**, Arg405 motion away from the active site. **G**, Asn403 twist away from active site.
